# Heritability and pre-adult survivorship costs of ectoparasite resistance in the naturally occurring *Drosophila*-*Gamasodes* mite system

**DOI:** 10.1101/2022.12.15.520609

**Authors:** Michal Polak, Joy Bose, Joshua B. Benoit, Harmanpreet Singh

**Author notes:** **Author Contributions:** M.P. designed the research, acquired funding for the research, contributed to conducting the research, analyzed data, wrote the manuscript; J.B. designed the research, conducted the research, analyzed data, contributed to editing the manuscript; J.B.B designed the research, acquired funding for the research, contributed to editing the manuscript; H.S. contributed to conducting the research, data organization and curation. **Declaration of interests:** The authors declare no competing interests. **Data accessibility:** Raw data will be made available on Mendeley Data.

## Abstract

Our understanding of the evolutionary significance of ectoparasites in natural communities is limited by a paucity of information concerning the mechanisms and heritability of resistance to this ubiquitous and diverse assemblage of organisms. Here, we report the results of artificial selection for increasing ectoparasite resistance in replicate lines of *Drosophila melanogaster* derived from a field-fresh population. Resistance, as ability to avoid infestation by naturally occurring *Gamasodes queenslandicus* mites, increased significantly in response to selection, and realized heritability (s.e.) was estimated to be 0.11 (0.0090). Ability to deploy energetically expensive bursts of flight from the substrate was a main mechanism of resistance that responded to selection, aligning with previously documented metabolic costs of fly behavioral defenses. Host body size, which affects parasitism rate in some fly-mite systems, was not shifted by selection. In contrast, resistant lines expressed significant reductions in larva-to-adult survivorship with increasing toxic (ammonia) stress, identifying an environmentally modulated pre-adult cost of resistance. Flies resistant to *G. queenslandicus* were also more resistant to a different mite, *Macrocheles subbadius*, suggesting that we documented genetic variation and a pleiotropic cost of broad-spectrum behavioral immunity against ectoparasites. This study demonstrates significant evolutionary potential of an ecologically important trait.

## Introduction

Parasites are ubiquitous in the environment (Price, 1980, Schmid-Hempel, 2021), and by definition, they damage host fitness (Goater et al., 2014). As a consequence, parasites can represent potent agents of natural selection, capable of driving the (co-)evolution of host defensive adaptations and altering host population genetic structure (Sheldon & Verhulst, 1996, Little, 2002). Acquiring estimates of genetic variation for host defensive traits is a key to predicting the evolutionary and ecological consequences of this selection (Wakelin, 1978, Endler, 1986, Henter & Via, 1995, Sorci et al., 1997).

There is now a large body of empirical evidence showing that heritable variation for resistance is indeed a common feature of natural populations of plant, animal and microbial species (Sorci et al., 1997, Hufbauer & Via, 1999, Lively & Dybdahl, 2000, Carius et al., 2001, Little, 2002, Duffy & Sivars-Becker, 2007, Foster et al., 2007, Schmid-Hempel, 2021). However, the existence of widespread heritability for resistance may also be considered paradoxical, as strong selection should act to deplete genetic variation at relevant loci (Barton & Turelli, 1989, Stearns, 1992). Several hypotheses for the maintenance of standing genetic variation for parasite resistance have been proposed, including frequency-dependent selection (Hamilton & Zuk, 1982, Lively & Dybdahl, 2000), and pleiotropic costs of resistance under heterogeneous parasite pressure (Parker, 1990, Rigby et al., 2002). Whereas the costs hypothesis has received considerable support, resistance costs have not always been found, fueling debate over their general importance (Simms & Rausher, 1987, Bergelson & Purrington, 1996, Coustau & Chevillon, 2000, Rigby et al., 2002). One proposed solution is that environmental variation can influence the expression of costs, although such effects also have been variable, and even contradictory in some cases (Coustau & Chevillon, 2000, Purrington, 2000, Sandland & Minchella, 2003, Osier & Lindroth, 2006, Boege et al., 2007, Cipollini et al., 2014). In addition to environmental influences, expression of costs may often depend on the stage of the life cycle, possibly reflecting shifting patterns of host resource allocation to different fitness functions across developmental stages (Sandland & Minchella, 2003, Boege & Marquis, 2005). For example, in birdsfoot trefoil, *Lotus corniculatus*, costs of producing defensive cyanogenic glycosides are highest during episodes of peak reproductive effort (Briggs & Schultz, 1990). A lack of detectable costs may thus only indicate that the appropriate ontogenetic stage and/or environmental conditions have not been considered (Sandland & Minchella, 2003).

Estimates of standing genetic variation for resistance and of associated costs in animal host-parasite systems are generally derived from studies of endoparasites—parasites that invade the host and reproduce within the body (Coustau & Chevillon, 2000). For example, we know a great deal about the mechanisms and magnitude of genetic variation for resistance to parasitoids attacking insects (Salt, 1970, Henter & Via, 1995, Carton & Nappi, 1997, Kraaijeveld & Godfray, 1997, Fellowes & Godfray, 2000, Carton & Nappi, 2001, Carton et al., 2005, 2008, Kim-Jo et al., 2019). In contrast, less information is available about the mechanisms and genetic bases of ectoparasite resistance (Gibson & Amoroso, 2022). Yet, ectoparasites, which attack the body surface of their hosts, are a diverse and ecologically important group of organisms (Behnke, 1990, Hopla et al., 1994, Clayton et al., 2010), and the fitness costs they impose on major life-history traits can be pronounced (Møller, 1990a, b, Forbes & Baker, 1991, Lehmann, 1993, Polak & Markow, 1995, Polak, 1996, Fitze et al., 2004). Host physiological impairment is a common outcome of heavy ectoparasite burdens through resource extraction, leading to anemia and significant weight loss, nutritional impairment and elevated risk of death and damaged reproduction (Nelson et al., 1975, Møller et al., 1990, Lehmann, 1993,). Ectoparasites may additionally serve as vectors of transmissible parasitic disease in plants and animals, including humans (Marshall, 1981, Behnke, 1990, Moran et al., 2008, Jaenike et al., 2007), and thus they are of enormous ecological, veterinary and medical importance, through both to the damage they inflict and the pathogens they transmit.

The behavioral mechanisms that animals may deploy against ectoparasites are diverse (Hart, 1990, Moore, 2002, Rigby et al., 2002, Thieltges & Poulin, 2008, Clayton et al., 2010, Hart & Hart, 2018). At the outset, behavioral defenses may serve to prevent contact and colonization of the host body, so-called “first-line” forms of defense (Leung et al., 2001, Schmid-Hempel & Ebert, 2003, Thieltges & Poulin, 2008, Poirotte et al., 2017). In addition to detection and direct avoidance mechanisms, first-line defenses may also involve avoiding certain habitats (Girard et al., 2021), feeding activities (Moore, 2002), nesting sites (Møller, 1990b), food patches (Anderson & McMullan, 2018), and conspecifics including potential mates (Borgia & Collis, 1989, Read, 1990, Kavaliers et al., 2003, Stephenson et al., 2018). After contact with parasites, hosts may reduce parasite burden by deploying brisk reflex movements directed at the parasites, self-grooming (e.g., foot scratching and rubbing), being groomed by others in the social group (allogrooming), and by self-medicating (Hart, 1994, Huffman, 1997, DeJoseph et al., 2002, Kupfer & Fessler, 2018, Ramanantsalama et al., 2018). Once established, ectoparasite feeding also generally elicits pronounced behavioral and immunological responses that may limit parasite feeding and development (Wakelin, 1996, Wikel & Alarcon-Chaidez, 2001, Owen et al., 2010). Thus, the defensive repertoire of hosts against ectoparasites spans a wide range of complex pre- and post-attachment mechanisms, all of which are in theory subject to selection pressure and prone to trade off against other fitness-related traits.

Here, we tested for heritable variation in first-line defenses in the host *Drosophila melanogaster*, as ability to avoid contact with mites and to prevent colonization of the body. These mechanisms serve to avoid the fitness costs of parasitism *per se*, which can be severe in *Drosophila*, occurring as reduced metabolic efficiency, fertility, mating success, and life span, and increased disease transmission (Polak, 1996, Jaenike et al., 2007, Polak et al., 2007, Polak & Markow, 1995, Greene, 2010, Horn et al., 2020). Mite attachment induces a suite of host immune and stress responses (Benoit et al., 2020), which collectively are likely to require non-trivial resource allocation (Lochmiller & Deerenberg, 2000). Flies also may repel attached mites by releasing free radicals and melanin at the feeding site (Lemaitre & Hoffmann, 2007, Lee & Miura, 2014), which are likely to pose additional physiological challenges and levy a fitness toll (Bian et al., 2010).

Our study system comprises the host *D. melanogaster* and its naturally co-occurring mite, *Gamasodes queenslandicus* (Acari: Parasitidae). These mites, which breach fly integument and consume host tissue while attached to their hosts (Polak and Spitz, manuscript; and see Polak, 1996), are generalist parasites recovered from a number of *Drosophila* species and other insects in Australasia (Halliday et al., 2005, Yao et al., 2020, M. Polak, personal observation). The fly’s “behavioral immunity repertoire” encompasses locomotor movements away from mites, and reflexive jumps and bursts of flight from the substrate, behaviors which generally are known to be metabolically expensive (Harrison & Roberts 2000, Zabala et al. 2009, Benoit et al. 2021). When a mite succeeds to grasp a fly, typically a tarsus, flies respond by vigorously prying and pushing at the mite and tarsal flicking to dislodge it (Polak 2003, Greene 2010). This suite of defensive behaviors closely matches those observed in the *Drosophila nigrospiracula-Macrocheles subbadius* mite association of the North American Sonoran Desert (Polak, 1996, Polak, 2003).

We employed artificial selection for increased pre-attachment host defenses and calculated realized heritability of mite resistance separately for male and female flies. Post-selection, a wing removal experiment was conducted to evaluate the importance of flight-related behavior in mediating the observed response to selection. Next, adult body size and larva-to-adult survivorship were contrasted between selected and control lines reared under conditions of increasing concentrations of ammonia (an environmental toxin), to evaluate whether any correlated evolutionary shifts occurred in these traits (Kraaijeveld & Godfray, 1997, Luong & Polak, 2007a). Host body size is an important fitness-related trait (Partridge et al., 1987, Flatt, 2020), and which mediates mite attachment in some drosophilid species (Campbell & Luong, 2016, Horn et al., 2020). Ammonia, a byproduct of metabolism, is an ecologically relevant stress factor known to accumulate to significant levels in larval substrates and to negatively influence the expression of fly life-history traits (Borash et al. 1998, 2000). Finally, we tested for cross-resistance in the selected lines to a different mite, *Macrocheles subbadius* (Acari: Macrochelidae), a cosmopolitan species known to parasitize drosophilid flies and also other insects (Polak & Markow, 1995, Polak, 1996); the aim here was to evaluate whether resistance to our focal mite could be generalized to another naturally occurring ectoparasite, and hence, whether the defensive traits we identified could have broader ecological significance.

## Materials and Methods

### Fly and mite laboratory populations

A laboratory base population of *D. melanogaster* Meigen was established with approximately 150 field-caught females and an equal number of males collected in February 2014 at two field sites 12 km apart (16° 5’4.59“S, 145°27’46.40“E; 16°12’50.19“S, 145°24’16.99“E) at Cape Tribulation, Queensland, Australia. Adult flies were collected directly from fruit substrates at both locations. Flies were returned to the laboratory and cultured on standard cornmeal-agar food media and controlled environmental conditions (12 h light (24°C): 12 h dark (22°C) in an environmental chamber. *Gamasodes queenslandicus* Halliday and Walter mites were harvested directly from the bodies of flies collected at both sites, and placed into culture medium to establish a large laboratory population (Polak 2003). The culture medium consists of a rich organic mixture of wheat bran, wood shavings, inactive yeast and bacteriophagic nematodes as a food source for the mites.

### Artificial selection for increased ectoparasite resistance

Artificial selection on *D. melanogaster* for increased resistance to *G. queenslandicus* mites was applied for 16 generations in each of three replicate fly lines, each independently derived from the base population described above. Each selected line had its own paired control line that was seeded each new generation with the same number of flies as its selected line. Flies used to seed each new generation of a given control line were unselected flies, randomly chosen from within their respective line that had not been exposed to mites. Selection lines are referred to as S1–S3, and control lines as C1–C3.

The protocol used to select for increased ectoparasite resistance is described in detail elsewhere (Polak 2003). Briefly, at each generation of selection, only male flies were exposed to mites in infestation chambers, composed of 500 mL glass jars containing ≈ 50 mL mite culture medium. Jars were sealed with breathable mesh to allow ample air exchange. For each selected line, 90 male flies were placed into each of 4 infestation chambers; 360 flies of each selection line were thus exposed to mites every generation. Flies interact with freely moving mites in these chambers on the surface of the medium, and parasitism occurs as in the field, with mites approaching and contacting flies from the substrate. Flies avoid contact with mites using a suite of distinct behaviors, and grooming to rid themselves of mites that have made contact (see Introduction). To apply selection, flies and mites were allowed to interact for 6-12 h in chambers until approximately 1/2 – 2/3 of flies acquired mites. All flies were recovered from chambers with an aspirator, and sorted under a stereomicroscope while anesthetized with a light stream of humidified CO_2_, and all flies with attached mites or mite-induced scars were discarded. Selection was applied by seeding each new generation of a given selection line with the unparasitized male flies. The proportions of selected flies (median, range) at each generation that were unparasitized after exposure are as follows: Line 1: 0.439, 0.292–0.79; Line 2: 0.444, 0.217–0.627; Line 3: 0.445, 0.314–0.79. Females were not selected, and were chosen at random from within their respective lines and paired (as virgins) with selected males to seed each new generation. The number of females used was 120 per line; 30 virgin females in each of 4 culture bottles per line. Selected males were distributed equally among these 4 bottles per line.

### Response to selection and realized heritability

To track response to selection, resistance assays were conducted at 6 time points over the course of the 16 generations of selection, following (Polak 2003; Luong and Polak 2007b). Each assay involved measuring resistance to ectoparasitism in each selected line relative to its paired control (unselected) line across a set of replicate infestation chambers. From 4–12 replicate chambers per pair of lines were used in each assay; the sexes were exposed separately. For a given pair of lines, groups of selected and control flies exactly equal in size were aspirated into a given chamber (total number of flies in each chamber ranged between 50 and 70 flies). Groups were distinguished by minute wing clips to the tip of one of the wings (≤ 3% of the wing) which themselves do not affect susceptibility to mites (Polak 2003; Benoit et al. 2020). Clips were administered to flies either to the right or left wing under CO_2_ anesthesia. The wing receiving a clip was alternated between the groups across replicate chambers. Flies were given a minimum of 24 h to recover from the clipping procedure prior to loading them into infestation chambers.

Flies and mites were allowed to interact in chambers for 6-12 h, until approximately 50% of flies were infested. Flies were recovered from chambers, scored for the presence of mites and scars, and identified as to their group of origin by their wing clips. Prevalence of parasitism in each group was calculated as the total number of infested plus scarred flies divided by the total number in each group that had been loaded into the chamber. Resistance was modeled as a threshold trait, with an expected underlying continuous variable called the liability, influenced by both genetic and environmental factors (Falconer and Mackay 1996). It was assumed that a single threshold separates resistant and susceptible forms (Polak 2003). Thus, mean prevalence across chambers of a given assay was transformed to “mean liability” for each treatment category (Falconer and Mackay 1996, p. 301), and the difference in mean liability (in standard deviation (SD) units) between selected and control groups was used to estimate the genetic improvement made by selection. To evaluate change in divergence over the 16 generations of selection, we used a generalized linear mixed model (GLMM) for each sex separately, with the following terms: replicate chamber, generation, selection treatment (i.e., selected vs. control), and the generation x selection treatment interaction; replicate line was nested within selection treatment and treated as a random effect. The generation x treatment interaction was of primary interest because it captured the effect of selection over time. The response variable was number of uninfested flies out of the total number of flies exposed to mites in a given group. Models were implemented in SAS (vers. 9.4) using the GLIMMIX platform, which uses the binomial distribution as the response distribution, allows random nested effects, and reports *F* statistics for tests of significance. Realized heritability, in turn, was calculated from the slope of the regression of divergence on generation number (Muir, 1986). Selection was applied to males only, so realized heritability was calculated as twice the slope for each regression (Roff 1997, p. 40). Heritability was estimated using data from these first 16 generations of selection. After the 16th generation, selection was applied every 2-3 generations to maintain divergence.

### Wing removal experiment

We tested the consequences of wing removal for resistance divergence between selected and unselected (control) flies, at two (consecutive) generations post-selection. The wings of each fly were removed by cutting each wing off at its base with ultra-fine surgical scissors. Wingless selected and unselected flies were exposed to mites in common infestation chambers, as above; replicate chambers were used for each pair of lines. To distinguish selected and unselected groups, we clipped the scutellar bristles of one of the two groups of flies. Whether the scutellar bristles were clipped was alternated between the groups across replicate chambers. Wingless flies were allowed to recover post-surgery for at least 24 h, and exposed to *Gamasodes* mites. Flies were removed from a given chamber when 40–60% of flies were estimated to be infested. Recovered flies were scored for the presence/absence of parasitism under CO_2_ anesthesia. Assays comparing selected and unselected flies with intact wings were conducted in parallel to verify divergence in resistance between these groups. In these assays, the wings of both groups were contacted with the scissors while under CO_2_ anesthesia but the wings were not removed. As for the wingless flies, selected and unselected flies were distinguished by scutellar bristle clips. Data were analyzed using generalized linear models in JMP® Pro (vers. 15.0.0), where the number of infested flies was the numerator and the total number of flies the denominator. Generation and selection treatment were factors in each model. Models were fitted separately by sex and wing treatment. The main objective was to assess degree of divergence between selected and unselected groups among flies with or without wings, by sex.

### Body size and pre-adult survivorship

The effects of selection for resistance on host body size, estimated as thorax length (Robertson & Reeve, 1952, Partridge & Fowler, 1992,), was tested at two time points. The first was immediately after generation 16, the generation at which sustained artificial selection was terminated and the realized heritability estimated. The second time point was after generation 21 of selection. At this second time point, the design was expanded to incorporate exposure to ammonia as a source of toxic stress (see below), to test for a possible interaction between selection treatment and environmental stress on body size. At the first time point, virgin females were paired with an equal number of males from within their respective line, and allowed to lay eggs on grape juice-agar petri dishes (10 cm diameter). Dishes with eggs were incubated at 25°C for 24 h, after which exactly 60 first-instar larvae were transferred with blunted dissection probes to food vials containing 6.0 mL standard cornmeal-agar medium. There were 2 vials for each of the 6 lines (3 selected and 3 control). Food vials with larvae were returned to incubator conditions, and flies were allowed to develop to adulthood. Ample pupations sites were provided to avoid larval drowning. Upon emergence, adult flies were allowed to age for 12 h and thorax length of flies measured using an ocular micrometer of a stereomicroscope. Thorax length was taken as the linear distance from the frontal edge of the thorax to the tip of the scutellum to estimate body sizes of both males and females (Robertson & Reeve, 1952). At the second time point, first instar larvae were harvested as above, and transferred to food vials with 6.0 mL medium comprising one of 4 concentrations of ammonium chloride (NH_4_Cl, Sigma): 0 g/L (0 M), 15 g/L (0.28 M), 20 g/L (0.374 M), and 25 g/L (0.47 M), bracketing the concentration range shown in previous work to impair homeostasis, and to exert effects of fly life-history traits, including larva-to-adult viability (Borash et al. 1998, 2000). There were 4 replicate vials per concentration. All emerging adults from experimental vials were harvested, counted, and preserved in 70% alcohol. Thorax lengths of a random subset of 5 male flies from each replicate vial were measured, as above. The effect of selection treatment on thorax length at both time points was evaluated with a restricted maximum likelihood (REML) mixed model, where replicate line was nested within selection treatment and treated as a random effect. Survivorship data were on a binomial scale and analyzed using GLMM, where the number of adult flies that emerged from a given vial was the numerator and the total number of flies that seeded each vial (i.e., 60) the denominator. Line was treated as a random effect, nested within selection treatment, as above.

### Cross-resistance to *Macrocheles subbadius*

To test for cross-resistance to *Macrocheles* mites in the lines selected against *Gamasodes* mites, resistance assays were conducted as described above, except that the mites to which flies were exposed in experimental infestation chambers were *M. subbadius*. Cross-resistance assays were conducted after 2 generations of relaxed selection under mass culture, following the 16 generations of artificial selection. Resistance to *Gamasodes* was assayed in parallel, so that degree of resistance to the mites could be directly compared. The mass culture of *M. subbadius* was established in the laboratory with mites harvested from flies (*D. nigrospiracula*) collected at necrotic saguaro cacti in Arizona (USA), at two sites: 33°20’43.05“N, 111°25’21.47“W, 33°21’42.52“N, 111°23’43.31“W. Mites were recovered directly from the bodies of field-caught *D. nigrospiracula* of both sexes under CO_2_ anesthesia, and cultured (Polak 2003). Data were analyzed using GLIMMIX in SAS (vers. 9.4) with generalized linear modeling, to test for difference between each resistant line against its paired control. Confidence intervals were obtained using JMP® Pro (vers. 15.0.0).

## Results

Artificial selection for increased resistance to *Gamasodes* mites applied for 16 generations to a field-fresh population of *D. melanogaster* resulted in strong evolutionary responses, reflected in a significant generation x selection treatment interaction on the probability of parasitism in both sexes (Table 1). The degree of divergence between selected and unselected lines increased steadily over the course of selection, and responses were congruent between replicate selection lines (Fig. 1A and B). Mean (s.e.) heritability estimates calculated from regression slopes for males and females separately, were 0.106 (0.00219) and 0.108 (0.0199), respectively (Table 2). Thus, the host population we sampled at Cape Tribulation, Australia, harbors significant additive genetic variation for resistance to *Gamasodes* mites. The notably steady and progressive evolutionary response suggests multifactorial inheritance, and the observation that responses were similar in magnitude between the sexes, despite selection having been applied to males only, implies autosomal effects.

**Figure 1.**
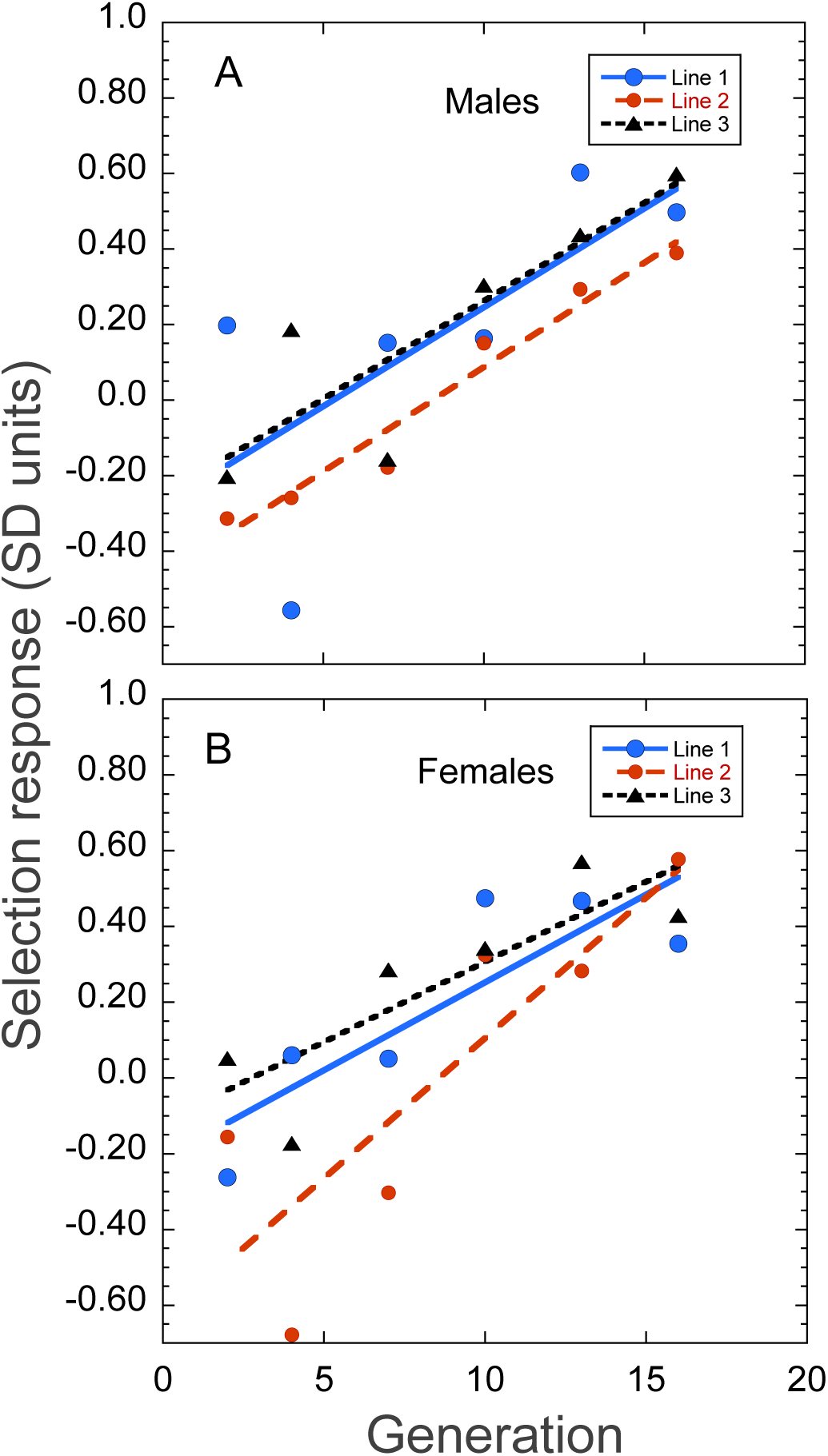
Evolutionary response of ectoparasite resistance in standard deviation (SD) unites across 16 generations of artificial selection in *D. melanogaster*, separately by males (A) and females (B). Selection was applied to male flies only, and response was tracked in both sexes. Table 2 provides the realized heritability estimates.

**Table 1.** Results of generalized linear mixed models on ectoparasite resistance in D. melanogaster, separately by sex. Selection on males for increasing resistance against G. queenslandicus mites was applied for 16 generations in three replicate host lines derived from a common base population. The significant generation x treatment interactions capture the significant responses to selection over the course of selection for both sexes.

**Males**
| Effect | Numerator<br>d.f. | Denominator<br>d.f. | *F* | *P* |
| --- | --- | --- | --- | --- |
| Chamber | 16 | 213 | 24.79 | < 0.0001 |
| Generation | 6 | 213 | 44.33 | < 0.0001 |
| Selection treatment | 1 | 4 | 1.92 | 0.238 |
| Generation x Selection trt. | 6 | 213 | 11.48 | < 0.0001 |

**Females**
| Effect | Numerator<br>d.f. | Denominator<br>d.f. | *F* | *P* |
| --- | --- | --- | --- | --- |
| Chamber | 16 | 211 | 15.40 | < 0.0001 |
| Generation | 6 | 211 | 39.33 | < 0.0001 |
| Selection treatment | 1 | 4 | 1.68 | 0.2649 |
| Generation x Selection trt. | 6 | 213 | 9.29 | < 0.0001 |

Line (nested within selection treatment) parameter estimate: 0.04263 (s.e., 0.0333)

**Table 2.**
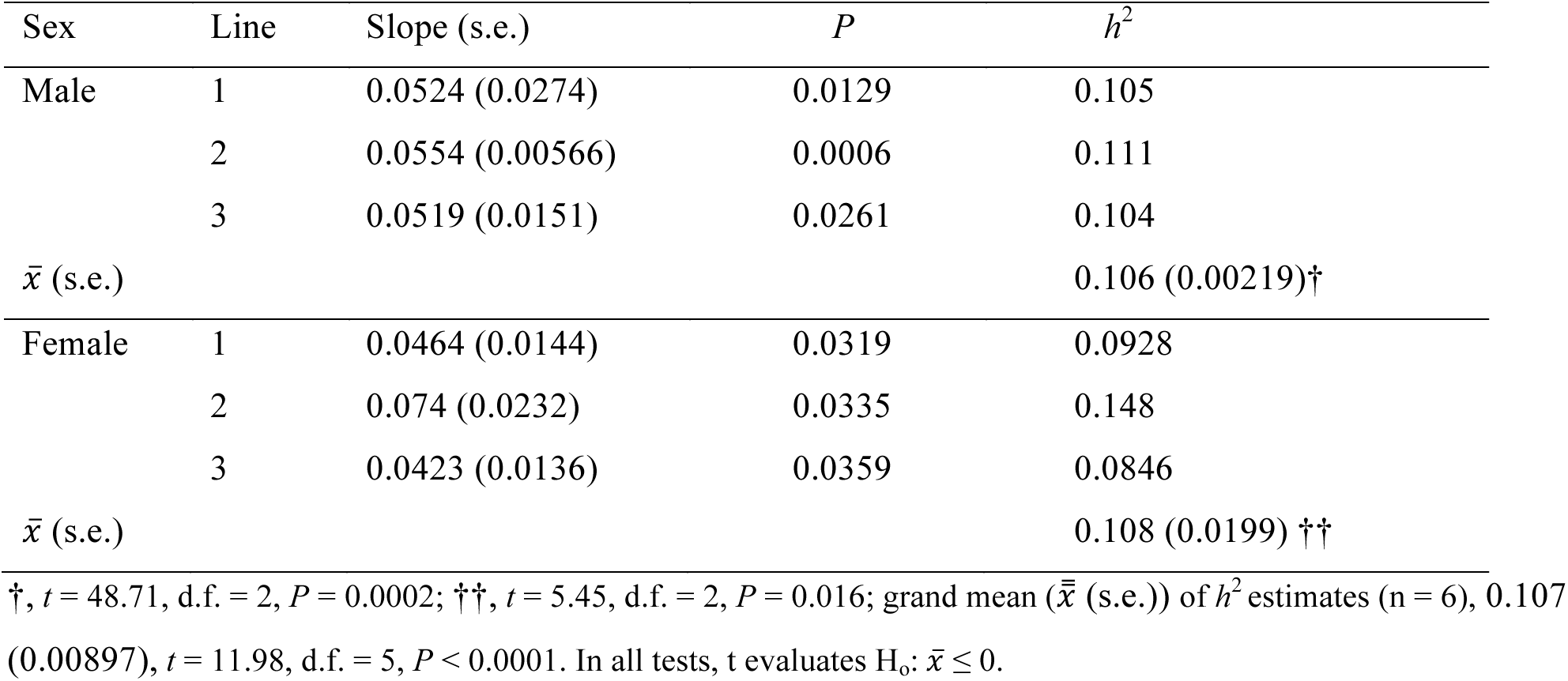
Realized heritability estimates of resistance in D. melanogaster to Gamasodes mites. Estimates are reported separately by sex and replicate selection experiment.

Among flies with experimentally removed wings, the effect of selection treatment on probability of parasitism was not significant for either males or females (Table 3A and B, respectively): selected lines exhibited similar levels of resistance compared to unselected flies, indicating that wing removal essentially eliminated the defensive advantage enjoyed by selected lines (Fig. 2). Parallel assays conducted at the same time with intact flies confirmed significant separation in resistance between selected and control lines, as expected (Table 3; Fig. 2). These results identify flight-related behaviors as a key component of ectoparasite resistance that responded to artificial selection.

**Figure 2.**
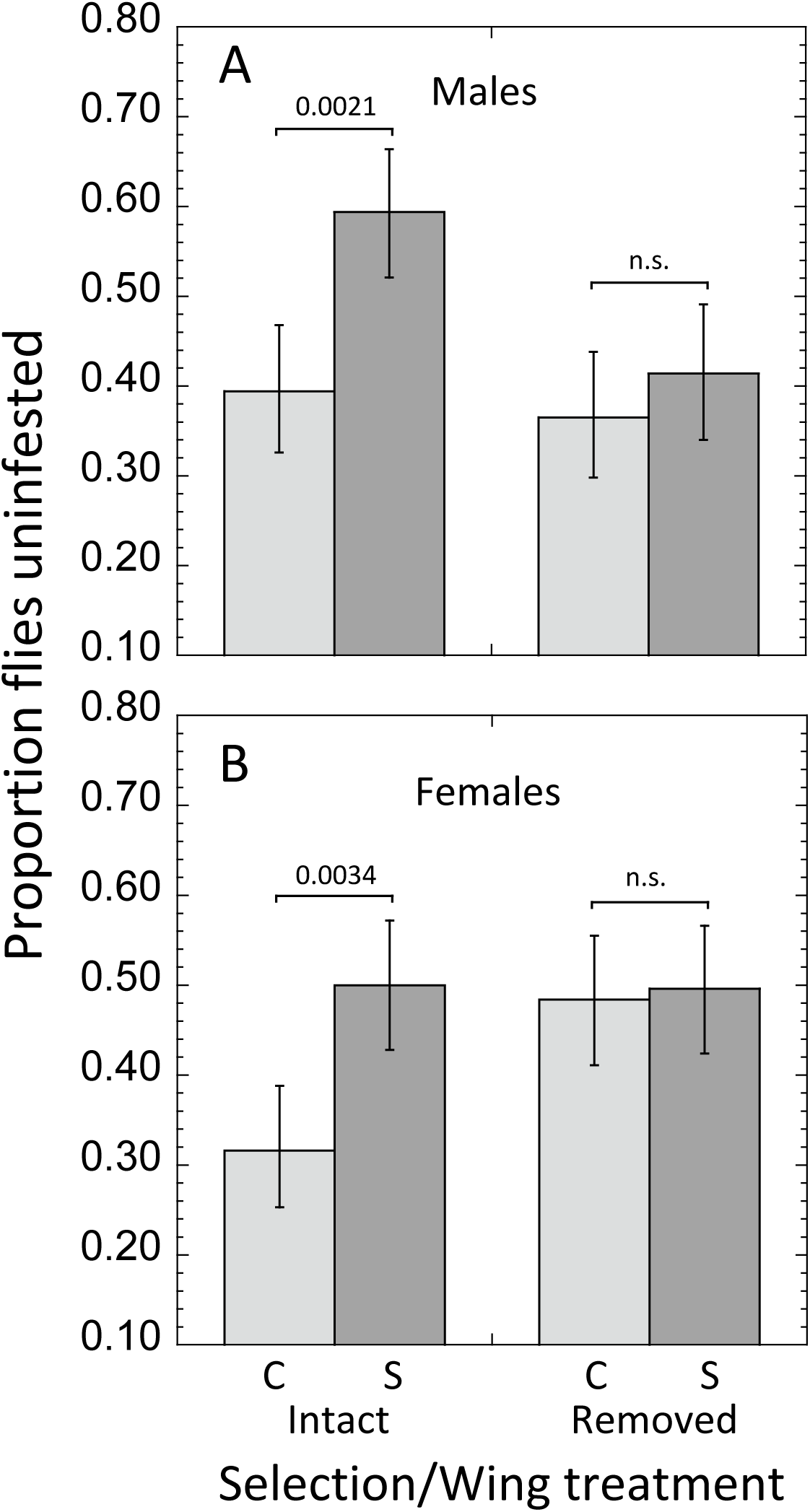
Results of the wing removal experiment for male (A) and female (B) flies. For both sexes, selected flies were more resistant than controls. For wingless flies, the separation in resistance between selected and control flies disappeared. Error bars represent 95% CIs.

**Figure 3.**
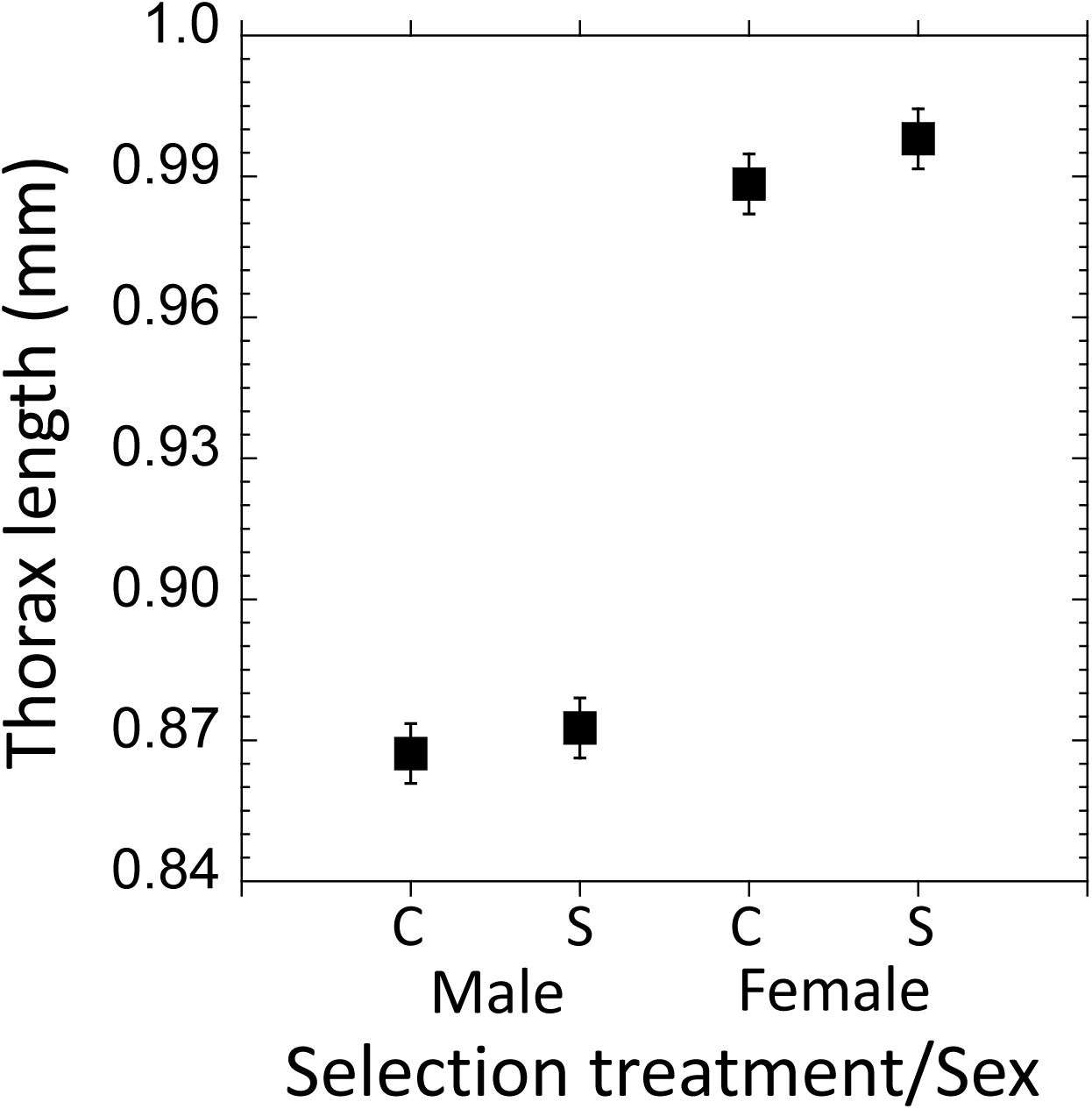
Mean (± 1 s.e.) thorax length for control and selected flies, by sex, following 16 generations of artificial selection for increased resistance against *Gamasodes* mites.

**Table 3.** Results of generalized linear models on ectoparasite resistance by wing removal treatment for (A) males and (B) females. For each sex, the effect of selection treatment on infestation probability was lost among wingless flies. Resistance assays conducted in parallel using intact flies exhibited significant selection treatment effects, confirming the significant divergence between selected and unselected flies, as had been documented immediately at the terminus of artificial selection.

| (A) Males: |  | Wings removed |  |
| --- | --- | --- | --- |
| Effect | d.f. | Chi-Square | *P* |
| Generation | 1 | 7.487 | 0.0062 |
| Selection treatment | 1 | 0.506 | 0.48 |
| Generation x Selection trt. | 1 | 0.104 | 0.75 |
|  |  | Wings intact |  |
| Effect | d.f. | Chi-Square | *P* |
| Generation | 1 | 9.352 | 0.0022 |
| Selection treatment | 1 | 14.130 | <0.001 |
| Generation x Selection trt. | 1 | 0.387 | 0.53 |
| *d.f., degrees of freedom |  |  |  |
| (B) Females: |  | Wings removed |  |
| Effect | d.f. | Chi-Square | *P* |
| Generation | 1 | 0.521 | 0.47 |
| Selection treatment | 1 | 0.0027 | 0.96 |
| Generation x Selection trt. | 1 | 0.284 | 0.59 |
|  |  | Wings intact |  |
| Effect | d.f. | Chi-Square | *P* |
| Generation | 1 | 0.930 | 0.93 |
| Selection treatment | 1 | 13.071 | <0.001 |
| Generation x Selection trt. | 1 | 0.519 | 0.47 |

Flies cultured under controlled larval densities post-selection did not show a significant difference in thorax length between selected and control lines, either for males or females (Table 4A). In a second iteration of this experiment (after 21 generations of selection), we evaluated both the effects of selection and ammonia treatments on adult (male) body size and egg-to-adult survivorship. For male body size, we again detected no significant effects of either factor, or of an interaction between them (Table 4B). In contrast, there was a strong overall effect of ammonia treatment on egg-to-adult survivorship (Table 5), which decreased sharply with increasing ammonia concentration (and see Borash et al. 1998, 2000). No significant effect of selection treatment on this trait was found (Table 5). Importantly, we detected a significant *interaction* between selection treatment and ammonia concentration on egg-to-adult survivorship (Table 5). Whereas selected and control lines did not differ at 0 g/L ammonia, selected lines showed progressively decreased survivorship relative to unselected lines with increasing ammonia concentration, from a 9.5% reduction at low concentration (15 g/L) to a 39.5% reduction at highest concentration (25 g/L) (Fig. 4). These results indicate an environmentally modulated pre-adult survivorship cost of ectoparasite resistance.

**Figure 4.**
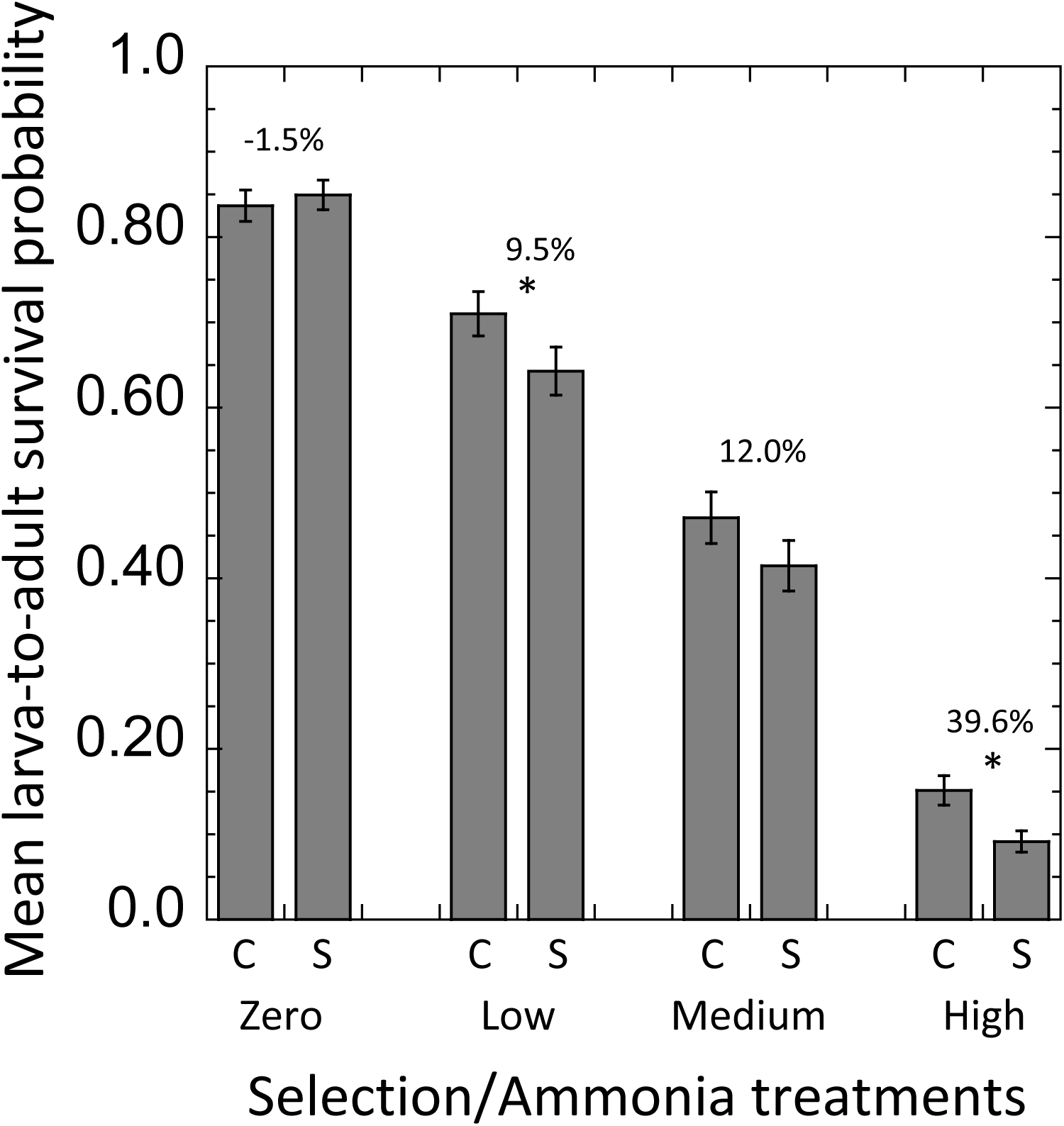
Probability of larva-to-adult survivorship across artificial selection and ammonia treatments. Survivorship among selected lines was significantly reduced with increasing ammonia present in the larval food substrate, reflected in a significant selection treatment by ammonia interaction (*P* = 0.003; Table 6). Error bars are +/-1 SE. Asterisks indicate a significant difference between selection treatments. Percent reduction in survivorship of selected lines relative to control lines is provided above each pair of means.

**Table 4.** Results of REML mixed models on body size in D. melanogaster following (A) 16 and (B) 21 generations of artificial selection for increasing resistance to G. queenslandicus mites.

(A) Factors in this experiment are sex, selection treatment (selected and control), and line.
|  | Numerator | Denominator | *F* | *P* |
| --- | --- | --- | --- | --- |
| Fixed effect | d.f. | d.f. |  |  |
| Sex | 1 | 232 | 583.460 | < 0.0001 |
| Selection treatment | 1 | 4 | 0.973 | 0.340 |
| Sex x Selection trt. | 1 | 232 | 0.137 | 0.712 |
| Line (nested within treatment and treated as a random effect) variance component: $4.44 \times 10^{-5}$ (s.e., $5.91 \times 10^{-5}$), $P = 0.452$. | | | | |

|  | Numerator | Denominator | *F* | *P* |
| --- | --- | --- | --- | --- |
| Fixed effect | d.f. | d.f. |  |  |
| Selection treatment | 1 | 4 | 2.536 | 0.186 |
| Ammonia treatment | 2 | 62 | 2.402 | 0.099 |
| Selection x ammonia trt. | 1 | 62 | 1.229 | 0.300 |
| Line (nested within treatment and treated as a random effect) variance component: $7.90 \times 10^{-5}$ (s.e., $1.41 \times 10^{-4}$), $P = 0.574$. | | | | |

**Table 5.** Results of a generalized linear mixed model on pre-adult (larva-to-adult) survivorship in lines of *D. melanogaster* selected for resistance to *G. queenslandicus* mites. Lines were reared at varying ammonium chloride concentrations (0, 15, 20, and 25 g/L) in the larval food. Line was treated as a random effect and nested within selection treatment.

| Effect | Numerator<br>d.f. | Denominator<br>d.f. | *F* | *P* |
| --- | --- | --- | --- | --- |
| Selection treatment | 1 | 4 | 2.52 | 0.187 |
| Ammonia treatment | 1 | 131 | 639.03 | <0.0001 |
| Selection x Ammonia trt. | 1 | 131 | 4.86 | 0.0031 |
| Line parameter estimate: 0.0335 (s.e., 0.0266) |  |  |  |  |

Flies selected for increased resistance to *G. queenslandicus* were strongly resistant to *G. queenslandicus*, and also to *M. subbadius* (Table 6; Fig. 5). Indeed, significant cross-resistance to *M. subbadius* was evident in all three replicate host lines selected for resistance to *G. queenslandicus* (Fig. 5), indicating a robust effect. Some degree of specificity of resistance to *G. queenslandicus* could nevertheless be discerned, as selected lines consistently exhibited relatively superior resistance to *Gamasodes* than to *Macrocheles*: selected lines 1–3 were 42.0%, 49.4% and 34.4% more resistant to *Gamasodes* relative to controls, and 26.6%, 32.4% and 25.8% more resistant to *Macrocheles*, respectively. Thus, the defensive mechanisms appear to be more effective against the ectoparasite used as the agent of selection.

**Figure 5.**
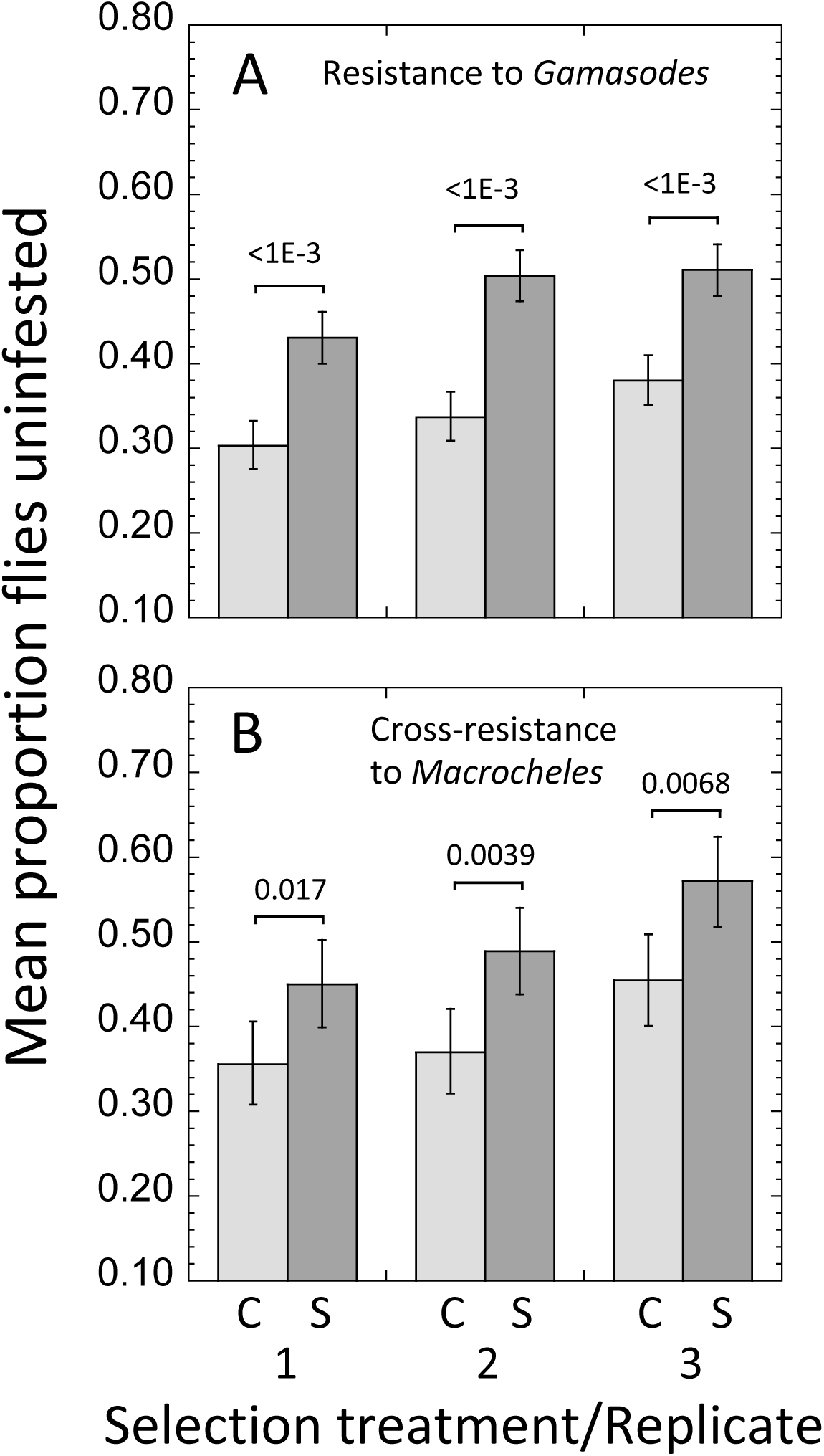
Resistance and cross-resistance in *D. melanogaster* after selection for increasing resistance to *G. queenslandicus* mites. Data are presented for control (C) and selected (S) host lines for each of the 3 replicates. Relative to controls, selected lines were strongly more resistant to *G. queenslandicus* (A), confirming the significant response to selection. Selected lines were also significantly cross-resistant to *M. subbadius* in all cases (B). Error bars are 95% CIs. Exact levels of statistical significance are from generalized linear modeling.

**Table 6.** Results of generalized linear models on ectoparasite resistance to Gamasodes (A) and Macrocheles (B) mites. Resistance assays were conducted following 16 generations of artificial selection for resistance to G. queenslandicus. Results indicate that selection lines consistently had superior resistance to G. queenslandicus, as expected, and to M. subbadius, indicating cross-resistance to this novel mite.

| (A) Resistance to *Gamasodes* |  |  |  |  |
| --- | --- | --- | --- | --- |
|  | Numerator | Denominator | *F* | *P* |
| Line | d.f. | d.f |  |  |
| 1 | 1 | 66 | 35.24 | < 0.0001 |
| 2 | 1 | 66 | 57.87 | < 0.0001 |
| 3 | 1 | 66 | 35.17 | < 0.0001 |

| (B) Cross-resistance to *Macrocheles* |  |  |  |  |
| --- | --- | --- | --- | --- |
|  | Numerator | Denominator | *F* | *P* |
| Line | d.f. | d.f |  |  |
| 1 | 1 | 22 | 6.65 | 0.0171 |
| 2 | 1 | 22 | 10.43 | 0.0039 |
| 3 | 1 | 20 | 9.10 | 0.0068 |

## Discussion

Trait heritability, the additive component of quantitative genetic variation, is a necessary condition for evolutionary response to natural selection (Endler, 1986). Here, we have documented significant heritability of ectoparasite resistance in a population of *D. melanogaster* recently derived from nature, demonstrating additive genetic variation for this trait; realized heritability (*h*^2^) of resistance was estimated to be 0.107, and values were consistent between the sexes, being 0.106 for males and 0.108 for females. Interestingly, the resistance to *Gamasodes* that we increased evolutionarily conferred cross-resistance to a different mite, *M. subbadius*, suggesting that we uncovered genetic variation for a general defensive ability against a given class of parasites (*sensu* Fellowes et al., 1999, Schmid-Hempel & Ebert, 2003). Furthermore, the defensive abilities we targeted could have functional significance in an even broader ecological context, as in interactions with predators such as ants (e.g., green tree ants, *Oecophylla*), which are a major class of natural enemy of flies observed at our field site, against which adult flies deploy similar evasive maneuvers as against mites (M. Polak, personal observation).

The heritability estimates we report are generally lower than for desert-endemic *D*. *nigrospiracula* (for which realized *h*^2^ of resistance to *M. subbadius* mites ranged from 0.12 to 0.15 (Polak, 2003, Luong & Polak, 2007b)), as well as for other host species (Møller, 1990a, Boulinier et al., 1997, Mazé-Guilmo et al., 2014, Yáñez et al., 2014, Buzatto et al., 2019). One possible explanation for the difference between the two *Drosophila* systems is that *Gamasodes* mites generate a more potent selection pressure than *Macrocheles*, which could have progressively depleted genetic variation in *D. melanogaster* over past generations. Moreover, the *Drosophila*-*Gamasodes* system we studied occurs in tropical Queensland (Australia), and predatory ants, as mentioned, are a notable source of mortality in the fly population (M. Polak, personal observation). Sustained pressure by multiple natural enemies on the same or overlapping set of defensive traits could have contributed to the depletion of host genetic variation for the defensive traits we studied.

Of further relevance is that *Gamasodes* mites are faster moving and aggressive, and more efficient at breaching host behavioral defenses than *M. subbadius*. The lower *h*^2^ of resistance to *Gamasodes* may also be the result of this higher parasite mobility, which would have the effect of reducing the number mite-fly interactions leading to attachment. Reducing the number of such events would increase the stochastic variance in the attachment rate, eroding the link between intrinsic (genetic) host factors and parasitism, hence reducing heritability (Hoffmann, 1999). A parallel argument has been made to explain variation in heritability of resistance to different ectoparasites attacking tropical cattle, where heritability estimates for numbers of ticks and buffalo flies were 0.34 and 0.06, respectively (Mackinnon et al., 1991). It was suggested that the sharply reduced *h*^2^ for fly counts could have been due to the high mobility of the attacking flies (Mackinnon et al., 1991).

Among flies whose wings were experimentally removed, selected flies lost their resistance advantage relative to their unselected counterparts. Wingless flies of both groups were able to deploy substrate-borne locomotor maneuvers such as running and jumping, but as a consequence of the manipulation, were prevented from deploying takeoff flights when approached or touched. The loss of resistance we observed indicates that the sensorimotor responses to mites we selected for require flight, most likely as microbursts from the substrate. An important aspect of this experiment is that both selected and unselected flies were wingless when they were exposed to mites. This design feature controlled for the possibility that the wing removal treatment altered fly attractiveness to mites, as wingless flies could have been perceived to be aberrant; mite choice in relation to host phenotype has been shown in other fly-mite systems (Campbell & Luong, 2016, Perez-Leanos et al., 2017, Horn et al., 2020). Thus, even if wing removal altered host attractiveness to mites, any such effect presumably would have affected selected and unselected flies similarly, minimizing confounding effects.

The discovery that flight-related behavior was central to the evolutionary response aligns well with previous physiological and transciptional analyses. It has been shown that flies that interacted with mites but that succeeded to avoid parasitism (i.e., the resistant class that we used for selection) exhibited reduced body condition, as decreased lipid and glycogen stores (Benoit et al. 2020). These findings are consistent with the fact that insect flight, particularly during takeoff, is energetically expensive and associated with metabolic expenditures several times that for resting and even running (Harrison & Roberts, 2000, Zabala et al., 2009). Within the experimental chambers we used for selection, fly responses to mites including bursts of flight are frequent and sustained, likely accounting for these observed negative effects on host nutrient reserves.

Previous work also has shown that flies that interacted with mites exhibited differential regulation of multiple genes associated with carbohydrate and lipid homeostasis (Benoit et al. 2020). These results identify potential candidate (metabolic) genes that may have underpinned the response to selection we documented here; indeed, there is ample evidence for genetic variation for metabolic genes related to flight parameters in *Drosophila* (Laurie-Ahlberg et al., 1985, Merritt et al., 2006). Neural factors underlying the escape response could also have been involved, for example, by affecting reaction times or fly “irritability”, as changes in transcript levels of several neurogenesis-affiliated genes were likewise noted (Benoit et al. 2020). Interestingly, select anti-microbial genes were increased, which may represent host anticipatory immune response to secondary infection by microorganisms (Kraaijeveld & Wertheim, 2009), should first-line defenses fail.

In the present study, we also tested for correlated responses to selection in both adult body size and pre-adult survivorship. We found that thorax length did not differ significantly between selected and control lines, either for males or females. From these results we conclude that the selection program did not significantly alter host body size in the present experiment (cf. Luong and Polak, 2007a). Given that body size can be a determinant of parasitism in other fly-mite systems (Campbell & Luong, 2016, Perez-Leanos et al., 2017, Horn et al., 2020), these results are noteworthy because they underscore the importance of flight-related performance *per se* as a major mechanistic determinant of the resistance phenotype.

Given the statistical invariance of body size, egg-to-adult survivorship was significantly decreased in the selected lines relative to controls and in complex fashion, as the strength of this effect was dependent upon ammonia exposure during development in a dose-dependent manner. In the absence of ammonia, selected and control lines did not differ significantly, but as ammonia concentration increased, resistant lines exhibited progressively reduced egg-to-adult survivorship relative to controls. These results demonstrate an environmentally modulated pleiotropic cost of resistance, and more generally provide support for the theoretical expectation that genetic correlations manifested in the form of trade-offs should often be affected by environmental conditions (Sandland & Minchella, 2003, Sgrò & Hoffmann, 2004). For example, stress factors, such as environmental toxins and temperature extremes, are likely to alter the effects specific genes have on different traits, and hence the expression of genetic correlations that link them (Sgrò & Hoffmann, 2004).

Importantly, a similar pattern of expression of this pre-adult cost has been documented in *D. nigrospiracula* selected for resistance to *M. subbadius* mites, a system endemic to the North American Sonoran Desert, which is a very different habitat from that which flies for the present study were derived, i.e., edges of rainforest habitats in northeastern Australia. In *D. nigrospiracula*, the larva-to-adult survivorship cost was also magnified under increasing stress, in this case with increasing larval density (Luong and Polak 2007a, and see (Kraaijeveld & Godfray, 1997)). Interestingly, the cost documented in *D. nigrospiracula* could therefore reflect exposure to the same environmental factor studied here, as ammonia, a byproduct of metabolism, builds up in larval substrates from larval excreta as densities rise (Borash et al. 1998, 2000). Overall, these results across different systems identify a potentially general cost of ectoparasite resistance expressed at the pre-adult stage of the host life cycle, likely to be sustaining genetic variation for a broad-spectrum adult resistance trait. Because our study system occurs naturally, and given that artificial selection was expressly applied to a field-fresh host population, it is concluded that ectoparasite resistance is an ecological important trait with significant evolutionary potential maintained in part by an environmentally modulated developmental cost of resistance.

## Acknowledgements

The research was supported by National Science Foundation (NSF) USA grant 1654417 (to M.P. and J.B.B.). We thank Rachel Black for her excellent assistance in media preparation.

## Notes

### Competing Interest Statement

The authors have declared no competing interest.

